# Habitat specificity is not universal proxy for sensitivity to habitat conversion among rodents on the Canadian Prairies

**DOI:** 10.1101/678268

**Authors:** Leanne M. Heisler, Gavin L. Simpson, Ray G. Poulin, Troy I. Wellicome, Britt D. Hall

## Abstract

Converting habitat for agricultural production threatens biodiversity loss worldwide and has significant implications for human well-being. Debates on how to conserve biodiversity as the demand for agriculture products rises is being informed by studies using habitat specificity as a proxy for sensitivity to land modification, assuming all species respond to habitat loss and fragmentation relative to their affinity towards the habitat type being converted. Here, we test this assumption among rodent assemblages on the Canadian Prairies, hypothesizing negative responses among grassland obligates and neutral or positive responses among habitat generalists to landscape change along a gradient of increasing agricultural intensity. We found complex, sometimes contradictory responses among rodent species, which depended on the magnitude of habitat loss that had occurred and did not always reflect each species’ relative affinity for grassland habitat. Our results suggest future studies should avoid assuming a single habitat type appropriately characterizes resource limitation among all species, and instead carefully consider which dimension of the ecological niche defines specificity for each species. Our results indicate habitat specificity is not always a reliable proxy for sensitivity to land modification, with significant implications for biodiversity conservation when used to inform agriculture and land use policies.

## Introduction

Agriculture is the leading cause of biodiversity loss worldwide, requiring the conversion of habitat into new land-use types for crop and livestock forage production, which reduces vegetation structure and simplifies composition (Benton et al. 2003; Connor et al. 2000; Tews et al. 2004; Fahrig et al. 2011). Loss of a single habitat patch results in species loss inside and outside the patch itself, reverberating as widespread biodiversity loss throughout the surrounding landscape (Burns & Grear 2008; Wilson et al. 2016). Over 75% of Earth’s terrestrial land mass shows evidence of land modification (Ellis and Ramunkutty 2008), of which 39-50% is due to modern agricultural and urban-industrial uses (Vitousek et al. 1997; Chapin et al. 2000; Foley et al. 2005). Globally, current rates of species extinction are 100-1000x higher than expected from the fossil record, indicating Earth’s 6^th^ mass extinction is underway (Tilman et al. 1994; Pimm et al. 1995; Barnosky et al. 2011; Dirzo et al. 2014). A crisis has also been identified in which ecological function is threatened in many biomes because of limited ecological protection (Hoekstra et al. 2005). As the global human population and food demand increases, further land modification will be needed, intensifying biodiversity loss and failing biome functions (Tilman et al. 2001; Foley et al. 2005; Green et al. 2005; Diaz et al. 2006; Tilman et al. 2011; Cardinale et al. 2012; Baudron and Giller 2013; Haddad et al. 2015).

Fragmentation effects on biodiversity are more complex, differentially influencing species’ habitat use, dispersal capabilities, and population persistence (Henle et al. 2004; Wilson et al. 2016). Fragmentation is a process by which remaining habitat patches become smaller and more isolated as land modification proceeds, changing the composition and spatial configuration of habitats at the landscape level (Fahrig 2003; Ewers and Didham 2005). Observed population responses to fragmentation are often inconsistent within and among taxonomic groups (Bender et al. 1998; Debinski and Holt 2000; Ewers and Didham 2005; Tylianakis et al. 2008; Fahrig 2017; Keinath et al. 2017). A recent meta-analysis found almost 75% of significant fragmentation effects indicate positive correlations with species abundances, further identifying the variable influence of habitat fragmentation (Fahrig 2017). Fragmentation is also highly correlated with habitat loss (Fahrig 2003); species may not respond until enough habitat loss occurs to make the change in landscape configuration relevant to the same species (Andren et al. 1994; Hanski and Ovaskeinan 2002; Betts et al. 2007). Population responses to one fragmentation metric may also be confounded with habitat loss or other fragmentation metrics. For example, area and edge effects can be both synergistic (Ewers and Didham 2007) and independent (Banks-Leite et al. 2010) but are often confounded because of their high collinearity (Fletcher et al. 2005) and scale dependency (Comfort et al. 2016). Population responses among species are also interdependent and indirect (e.g., competitive interactions, multi-trophic relationships, pathogen infection intensities, mutualisms between plants and animals; Tylianakis et al. 2008; Didham et al. 2012; Wilson et al. 2016). Despite all of this, recent assessments of fragmentation effects on biome-wide ecosystem function indicate the immediate need to better understand and act to reduce these effects where they negatively affect biodiversity (Haddad et al. 2015).

Land modification may homogenize plant and animal communities by favouring habitat generalists at the expense of habitat obligates. Habitat specificity is an evolutionary strategy for resource acquisition within heterogeneous landscapes. Species adapt to either acquire resources within one or a few habitat types and maintain resource availability through competition (i.e., habitat obligate), or use a diverse resource base across multiple habitat types and maintain access through colonization (i.e., habitat generalist; Futuyma and Moreno 1988; McKinney and Lockwood 1999; Marvier et al. 2004; DeVictor et al. 2008; DeVictor 2010; Clavel et al. 2011). Communities maintaining populations of both obligates and generalists are resilient to environmental change because species loss from some functional groups are counterbalanced by gains in other groups (McNaughton 1977; Chapin et al. 2000; Diaz et al. 2006). This also maintains ecosystem function by maintaining redundancy among groups experiencing loss during periods of environmental change (Chapin et al. 2000; Virginia and Wall 2001; Hector and Bagchi 2007). However, land modification permanently and rapidly homogenizes habitat mosaics into highly productive crop monocultures that experience consistent, intra-annual disturbances (Benton et al. 2003). Habitat generalists may be more capable of taking advantage of this high productivity because of their ability to colonize consistently disturbed habitats (Tilman et al. 1994; Wright et al. 2012). In contrast, habitat obligates are less flexible in the resource base they can use, and so are more vulnerable to disturbances that eliminate habitat (Tilman et al. 1994; DeVictor et al. 2008; Reino et al. 2014). The resulting generalist-dominated communities may synchronize assemblage responses to environmental change, with implications for ecosystem resilience (McKinney and Lockwood 1999; Chapin et al. 2000; Diaz et al. 2006; Clavel et al. 2011; Cardinale et al. 2012).

Here, we use rodent species responses to land modification to examine whether habitat specificity predicts sensitivity of rodent assemblages to habitat loss and fragmentation on the Canadian Prairies. Little is known regarding how rodent assemblages respond to the fragmentation of remaining habitat in agroecosystems, with implications for the ecosystem processes they maintain, including predator population regulation (Korpimäki et al. 2004; White et al. 2013), disease and parasite prevalence (Ostfeld and Holt 2004), plant diversity through foraging habits (Sieg 1987), soil conditions, nutrient flow, and habitat availability through burrowing activities (Grant and French 1980). Population abundance or density is typically estimated using conventional trapping, which has logistic constraints that limit studies to patch area and isolation effects (Hanser et al. 2011). Fewer studies have identified rodent species responses to fragmentation at the landscape level at which fragmentation affects population persistence (e.g., Pena et al. 2003; Heisler et al. 2013; Massa et al. 2013; Torre et al. 2015). The Canadian Prairies experienced spatially inconsistent land modification post-European settlement (Samson and Knopf 1994; Samson et al. 2004), resulting in landscapes that differ in the magnitude of grassland loss and fragmentation that occurred there (Gage et al. 2016), providing ideal conditions for this study.

We hypothesize that rodent species respond differently to fragmentation depending on the degree of habitat specificity each species exhibits towards grassland habitat. We predict grassland and vegetation obligates will respond negatively to increased loss and isolation of remaining grassland patches, while habitat generalists will either 1) show no response because individuals perceive no difference among habitat and land modified for agricultural use, or 2) exhibit positive responses to the introduction of highly productive, new land-use types and edge habitat providing access to additional resources.

## Methods

### Study area

Samples were collected across almost 2 million km^2^ of the mixed-grass prairie and aspen parkland of the Northern Great Plains in Canada (Fig. 1). Vegetation of the mixed-grass prairie consists mostly of grasses (e.g., *Agropyron* spp., *Bouteloua* spp., *Stipa* spp.), upland sedges (e.g., *Carex* spp.), forbs, club moss (e.g., *Selaginella densa)* and intermittent patches of small shrubs (e.g., *Symphoribarpus* spp., *Elaiagnus* spp., *Prunus* spp., *Amelanchier* spp.) and trees (e.g., *Populus* spp., *Salix* spp.) where moisture permits (Coupland 1961). The aspen parkland represents the transition zone between the southern mixed-grass prairie and northern boreal forest and is characterized by aspen bluffs (e.g., *Populus* spp.) and fescue prairie (e.g., *Stipa festuca;* Coupland 1961; Barker and Whitman 1988). Vegetation composition in the study area is heavily influenced by local topography, boulder clay soil deposits of varying texture, and a continental climate of extreme variation in temperature and precipitation with a short growing period of 3-5 months (Coupland 1950).

**Fig. 1.**
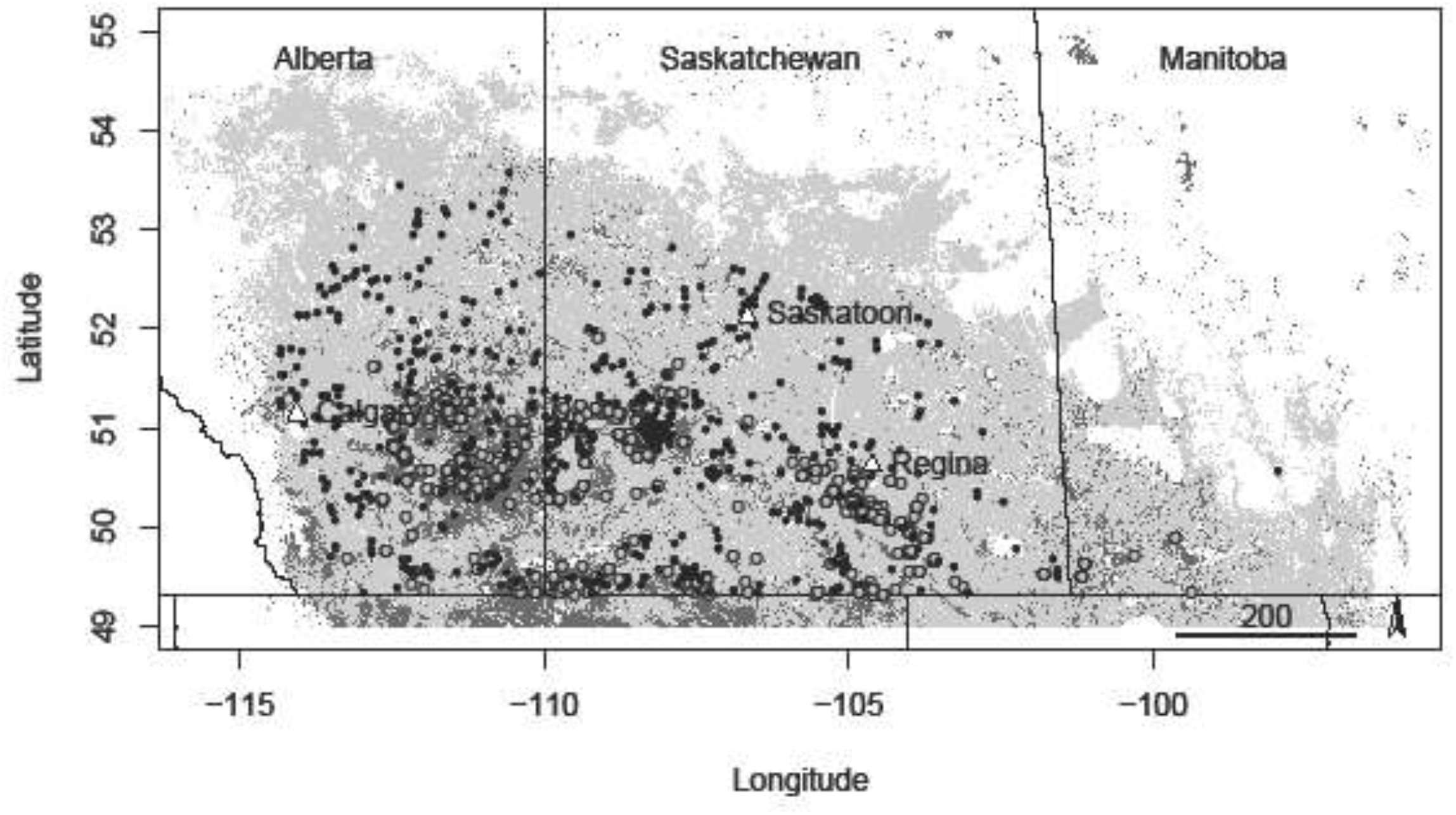
Distribution of owl pellet collection locations (black dots) in grasslands modified for agricultural use (cropland = shaded light grey; grassland = dark grey) in Alberta (left), Saskatchewan (middle), and Manitoba (right; lines = provincial boundaries) of Canada. Major cities in Alberta (i.e., Calgary and Edmonton) and Saskatchewan (i.e., Regina and Saskatoon) are given for reference. White areas represent waterbodies and regions outside the study area.

Both ecoregions are heavily altered by land modification, through which heterogeneous grasslands were converted into a few agricultural land use types, thus decreasing vegetative species composition and structure and increasing bare ground (Smoliak 1988). Only 6% and 27% of the aspen parkland and mixed-grass prairie remain intact, respectively (Roch and Jaeger 2014). Most modification occurred between the 1880’s and 1940’s (Smoliak 1988; Samson et al. 2004) but it continues today at lower rates. An average annual 2% loss has occurred since 2009 in some regions of the Canadian Prairies, resulting in an additional 13% loss of remaining grasslands (Gage et al. 2016). Land modification was not evenly distributed across the study area. Large tracts of grassland remain intact in southeast Alberta and southwest Saskatchewan, while over 90% of the grasslands in other regions are heavily modified (Roch and Jaeger 2014; Fig. 1). This variation in agricultural intensification provides an opportunity to examine landscape-level rodent species responses to grassland loss and fragmentation, with comparisons of responses among grassland obligates and habitat generalists.

### Rodent abundance estimates

Rodent species abundances were estimated from great horned owl *(Bubo virginianus*) and burrowing owl *(Athene cunicularia)* pellets collected across the study area (Heisler et al. 2016). Over the last 20 years, owl nests and associated roosts were visited once to several times during the breeding season and all accessible pellets were collected. Great horned owl pellets were collected from 207 locations by Alberta’s Environment & Sustainable Resource Development Fish & Wildlife division in 2000 and 2001, and from an additional 436 locations by research scientists and bird enthusiasts from 2008 until 2016. Burrowing owl pellets were collected from 1,179 locations by the Royal Saskatchewan Museum and the Canadian Wildlife Service from 1997 until 2016.

All pellets were processed to clean the bones and teeth in one of two ways: (1) soaked in 10% sodium hydroxide solution for 2 – 3 hours to dissolve the fur from the pellets; or (2) soaked in water and manually separated. Prey species were then identified by the diagnostic characteristics of craniomandibular elements (i.e., teeth, mandibles, skulls), using a reference collection provided by the Royal Saskatchewan Museum and Royal Alberta Museum. Species abundances were quantified for each collection based on the maximum number of right or left mandibles, or the total number of skulls present (i.e., minimum number of individuals). The resulting dataset includes 84,196 individuals of 11 grassland rodent species.

Owl pellets accumulated over differing amounts of time at each pellet collection location, requiring standardizing to account for differential sampling effort. Great horned owl pellet collection locations were visited once and likely represent owl foraging efforts for that year. In contrast, burrowing owl pellet collection locations were visited once to several times during the breeding season, and some locations were visited over multiple years. To account for these differences, rodent abundances were summed when more than 1 visit was made each year to the same pellet collection location. Each pellet sample therefore represents all pellets collected from a single location within a single year, hereafter referred to as a sample. To accurately reflect rodent community composition and be included in subsequent statistical analyses, each pellet sample also met a threshold of the minimum number of individuals identified necessary to reach the average prey species richness represented in pellet collections for both owl species (great horned owl, 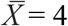 species; burrowing owl, 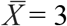 species). We therefore used great horned owl pellet samples with 30 or more individuals, and burrowing owl pellet samples with 10 or more individuals, respectively. A total 323 great horned owl and 1,061 burrowing owl samples were used in subsequent statistical analyses (Table 1).

**Table 1.**
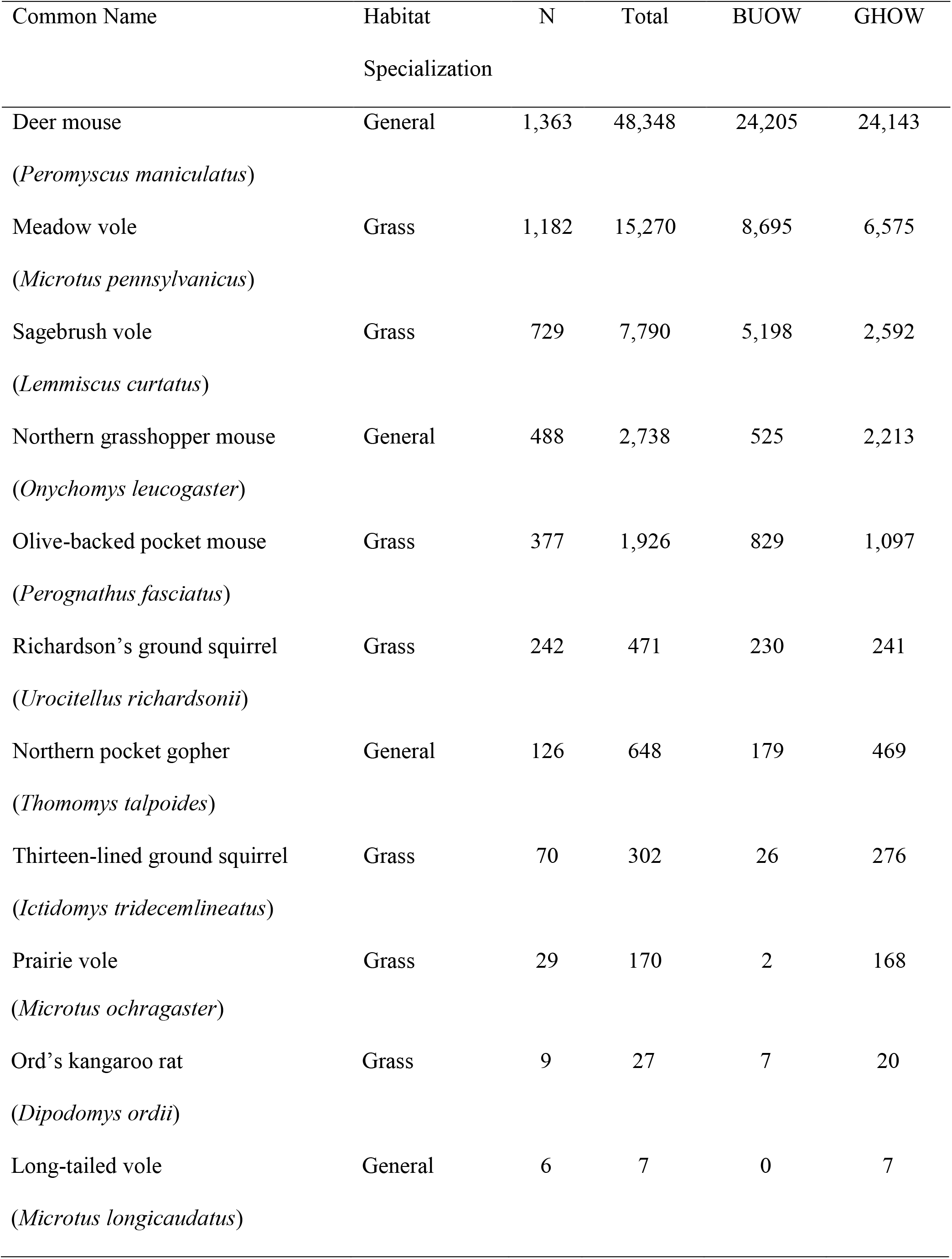
Frequency among samples (N) and abundance (Total) of rodent species estimated from great horned (GHOW) and burrowing owl (BUOW) pellet samples. Characterization as grassland obligates (Grass) or habitat generalists (General) based on habitat descriptions in Eder and Kennedy (2011).

Each rodent species was categorized as a grassland obligate or a habitat generalist according to its habitat description in Eder and Kennedy (2011; Table 1). Grassland habitats are dominated by herbaceous and shrub vegetation maintained by fire, grazing, drought, and freezing temperatures (Axelrod 1985; White et al. 2000). Vegetation composition and structure is largely determined locally by topography, precipitation, and soil texture (Epstein et al. 1997; Lane et al. 1998; Hooke and Burke 2000). Here, we use species natural history accounts from Eder and Kennedy (2011) to characterize grassland obligates as those species selecting for habitat vegetated with a combination of native grasses, forbs, and short woody plants, and habitat generalists as those species that occupy a variety of habitat types and are not limited by vegetation cover (McKinney and Lockwood 1999; Marvier et al. 2004; DeVictor et al. 2008; DeVictor et al. 2010; Clavel et al. 2011).

### Metrics of habitat loss, fragmentation, habitat composition, soil texture, and climate

We used several metrics to characterize habitat composition, grassland loss and fragmentation, and abiotic conditions surrounding pellet collection locations. Habitat composition (i.e., relative proportions of habitat types within landscapes), percent habitat loss, and several metrics of grassland fragmentation were quantified across the Canadian Prairies from raster thematic data generated from Landsat5-TM and Landsat 7-ETM+ multi-spectral imagery of 30 m resolution, representative of circa-2000 conditions (Agriculure & Agri-Food Canada 2012), which was reclassified to 6 broad land use (i.e., crop, tame forage, urban) and habitat categories (i.e., grass, woody, riparian).

Habitat composition was estimated using the proportion of each habitat type (i.e., grassland (GRASS), forest and shrub (WOOD), riparian (RIPARIAN)) and agricultural land-use type (i.e., cropland (CROP), tame forage (TAME), urban (URBAN)) within the foraging range surrounding each owl pellet collection location (Table 2). The influence of habitat loss was estimated using the cumulative percentage of the landscape comprised of crop, tame forage, and urban developments (LOSS; Table 2). Habitat fragmentation was characterized using measures of grassland edge density (m/ha; ED), average grassland patch perimeter-area ratio (i.e., shape complexity; PARA), average grassland patch size (ha; AREA), grassland patch density (number of patches per 100 ha; PD), and grassland patch cohesion (i.e., monotonic index of patch connectivity; COH; Table 2).

**Table 2.**
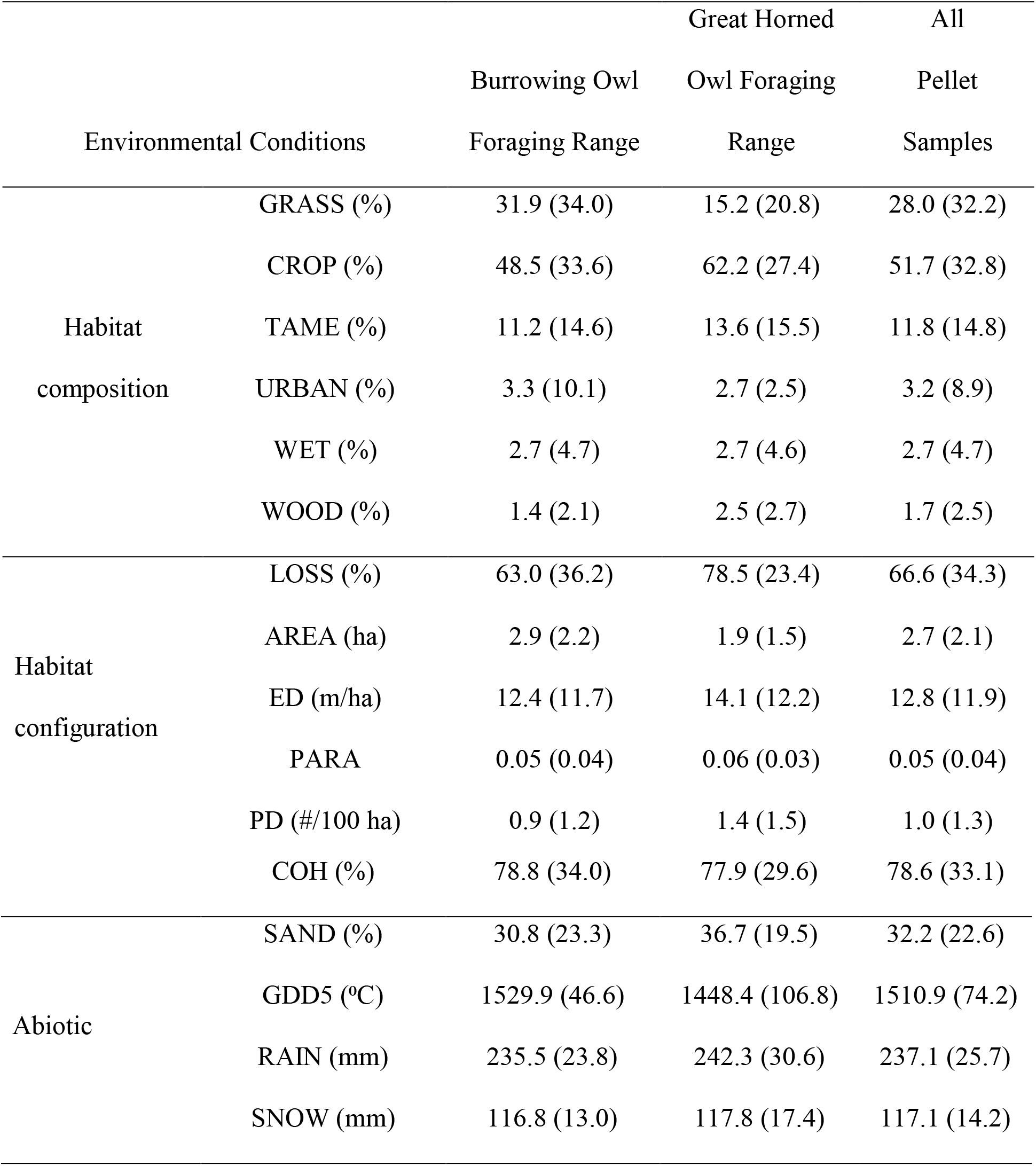
Average (+/− standard deviation) percent habitat loss (LOSS), habitat fragmentation (i.e., average grassland patch area (AREA), grassland edge density (ED), average grassland patch shape edge to area ratio (PARA), grassland patch density (PD), and grassland patch cohesion (COH)), habitat composition (cropland (CROP), grassland (GRASS), perennial livestock forage (TAME), urban development (URBAN), riparian areas (WET), and woody vegetation (WOOD), climate (total growing degrees >5 °C (GDD5), summer rainfall (RAIN), and winter snowfall (SNOW)), and soil texture (sand (SAND)) within a 2.5 km radius from each burrowing owl and great horned owl pellet collection location. Fragmentation metrics were taken from VanDerWal 2015.

We used other abiotic variables known to influence small mammal assemblage composition across the study area to produce more complete statistical representations of landscape-level habitat associations for each rodent species. We compiled soil texture variables from existing soil survey maps by Agriculture and Agri-Food Canada (Soil Landscapes of Canada Working Group 2010; Heisler et al. 2013). Soil texture was characterized from existing digitized soil survey maps using the mean proportion of sand (SAND) in the soils of each owl foraging range surrounding owl pellet collection locations (Centre for Land and Biological Resources Research 1996; Table 2).

We generated three climate variables from monthly precipitation and temperatures averaged over a 30-year period (i.e., 1971-2000; Agriculture and Agri-Food Canada 2013; Heisler et al. 2013; Heisler et al. 2014). Growing degree days were generated as the cumulative temperature above 5 °C from May to September averaged from 1971 to 2000 (GDD5), while the total annual precipitation from May to September and from October to April averaged from 1971 to 2000 characterized summer rainfall (RAIN) and winter snowfall (SNOW), respectively (Table 2). Lastly, we included habitat composition as the proportions of each land-use and habitat type (Heisler et al. 2013).

Metrics were calculated within radii varying in distance (i.e., 1 – 4 km at 0.5 km intervals) from pellet collection locations to estimate the appropriate scale of effect reflecting the foraging distances of burrowing owls and great horned owls from nests and associated roosts (Heisler et al. 2013). We used generalized additive models (GAMs) with a negative binomial distribution including the most abundant species (i.e., deer mouse) as the response and all fragmentation metrics as splines for each radius (see below for model structure). We chose a scale of effect with the least amount of information loss, or low Akaike Information Criterion (AIC), in both the burrowing owl and great horned owl GAMs so that all further statistical analysis could be done in a single GAM for each rodent species. Each metric was therefore estimated within a 2.5-km radius of each owl pellet collection location (Burnham and Anderson 2002).

All reclassifying, data preparation, and metric calculation was done using R version 3.5.3 (R Development Core Team 2019). The package raster (Hijmans 2019) was used to read and mosaic tiles, reclassify land cover, and prepare soil and climate data. Packages rgdal (Bivand et al. 2019) and raster were used to read and prepare owl pellet, soil, and climate data. Package landscapemetrics (Hesselbarth et al. 2019) was used to calculate landscape metrics.

### Statistical analyses

Non-linear responses of rodent species to fragmentation were identified using generalized additive models (GAMs). Land modification for agricultural use is a highly correlated, spatially explicit, sometimes non-linear process (Fahrig 2002) that may be reflected in rodent species responses to fragmentation. GAMs examine both linear and non-linear relationships using generalized regression of multiple predictors (Hasti and Tibshirani 1990; Wood 2006; Zuur et al. 2009; Zuur 2012). We fit GAMs for each species starting with a Poisson distribution and log link to account for positive integers:

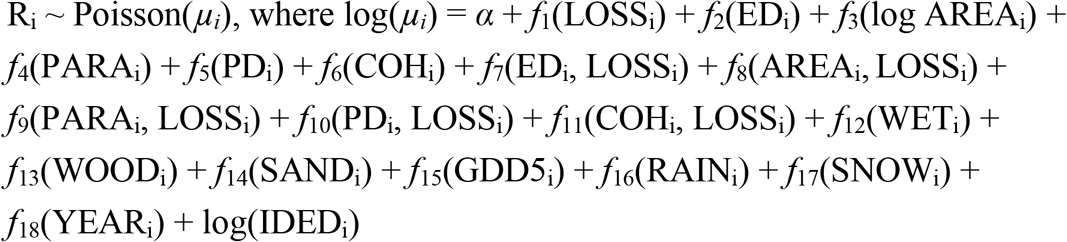

where R_i_ is rodent abundance in the i^th^ observation (pellet sample). The terms *f*_1_ – *f*_18_ are smooth functions of habitat loss (LOSS), grassland edge density (ED), average grassland patch size (AREA), average grassland patch shape complexity (PARA), grassland patch density (PD), grassland patch cohesion (COH), percent riparian (WET), percent woodland (WOOD), percent sand in soils (SAND), average annual growing degree days >5 °C (GDD5), average summer rainfall (RAIN), average winter snowfall (SNOW), and the year samples were collected (YEAR). The natural log-transformed sample size (IDED) was also included as an offset variable. The restricted maximum likelihood (REML) was used to avoid overfitting while estimating smoothing parameters. Parameters were penalized to zero and removed from the model during smoothing parameter estimation when they had little to no effect on rodent abundance. Overdispersion was accounted for by changing distributions to a negative binomial distribution when necessary (Zuur et al. 2009; Zuur 2012). Species abundance was included as the response. Species absences were included only when they were located within the species’ geographic range. To account for high concurvity between LOSS and fragmentation metrics (i.e., ED, AREA, PARA, PD, and COH), each fragmentation metric was included as a tensor product (i.e., synergistic effect) and tensor product interaction with LOSS (i.e., independent effect; Wood 2006). Habitat composition, and soil and annual climate conditions were included to improve model fit (Heisler et al. 2013; Heisler et al. 2014). Year was included to account for variable environmental conditions among years samples were collected. Total rodent individuals per sample was included as a log-transformed offset to account for differing sampling effort. Concurvity among all predictors was then assessed, those showing high concurvity with LOSS or fragmentation metrics were removed from the model (Ramsay et al. 2003).

Null models for each species included smooth functions of soil texture, annual climate conditions, habitat composition, year, and the offset term. Comparisons of the fitted and null models using AIC tested for statistical influence of habitat loss and fragmentation (Burnham and Anderson 2002). Only models >10 delta AIC units (i.e., competing models) from the null model were considered. The variance explained by fragmentation of each fitted model was estimated by subtracting the deviance explained of the null model from that of the fitted model. Inferences were made only from statistically significant predictors using p < 0.001 to account for variability in estimated p values (Zuur et al. 2009; Zuur 2012).

All statistical analyses were done using R version 3.6.3 (R Development Core Team). GAMs were estimated using the mgcv (Wood et al. 2016) and gamlss (Rigby and Stasinopoulos 2005) packages.

## Results

Six of the eleven rodent species considered here responded to metrics of habitat loss and fragmentation on the Canadian Prairies. Fitted models for four grassland obligates (i.e., meadow vole, prairie vole, Ord’s kangaroo rat, and Richardson’s ground squirrel) and one habitat generalist (i.e., long-tailed vole) were competitive with their corresponding null models, suggesting these models did no better at explaining variance in abundance than random chance (Table 3). Null models were not competitive for the remaining six species (Table 3), of which three were grassland obligates (i.e., sagebrush voles, olive-backed pocket mice, and thirteen-lined ground squirrels) and three habitat generalists (i.e., deer mice, northern grasshopper mice, and northern pocket gophers). Three species responded to the main effect of habitat loss (i.e., deer mice, olive-backed pocket mice, and northern pocket gophers; Table 4). All six species responded to at least one metric of fragmentation. Fitted models explained 35-82% of variance in abundance for each rodent species, of which fragmentation effects accounted for <11% for all species (Table 3).

**Table 3.**
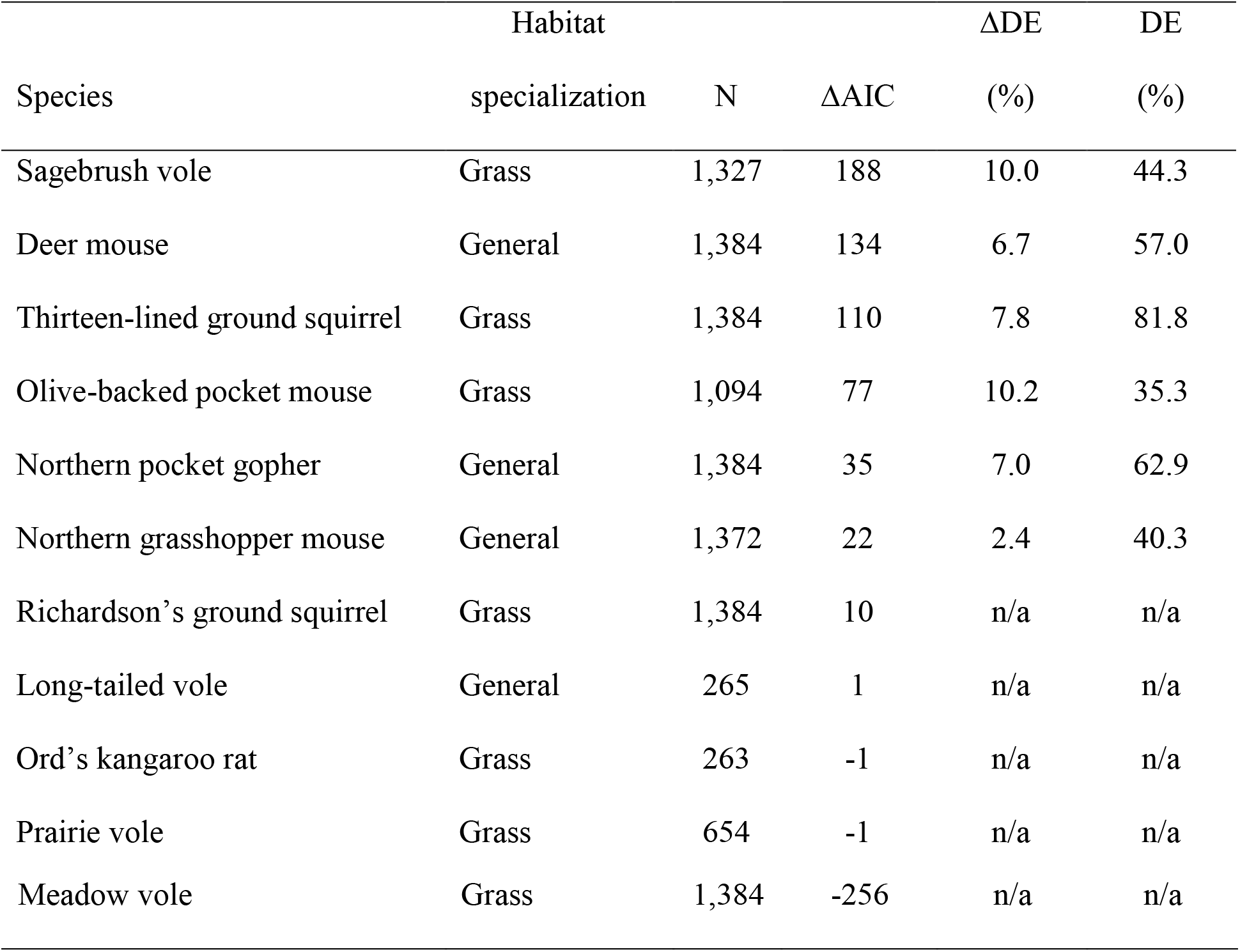
Model performance of fitted generalized additive models estimating change in abundance of each rodent species to habitat loss and fragmentation on the Canadian Prairies. The number of samples used to model each species’ response (N) differed among species because only samples within each species’ geographic range were used. All species models were fit using mgcv, except the deer mouse and sagebrush vole models, which were fit using gamlss. Statistical influence of each model was estimated by subtracting the Akaike Information Criterion (AIC) of each fitted model from the AIC of each corresponding null model (ΔAIC). Only models >10 ΔAIC units from the null model were considered. The strength of each fitted model was estimated by deviance explained (DE). The strength of habitat loss and fragmentation effects alone in each model was estimated by subtracting the DE of each null model from that of each corresponding fitted model (ΔDE).

| Species | Habitat | N | ΔAIC | ΔDE | DE |
| --- | --- | --- | --- | --- | --- |
|  | specialization |  |  | (%) | (%) |
| Sagebrush vole | Grass | 1,327 | 188 | 10.0 | 44.3 |
| Deer mouse | General | 1,384 | 134 | 6.7 | 57.0 |
| Thirteen-lined ground squirrel | Grass | 1,384 | 110 | 7.8 | 81.8 |
| Olive-backed pocket mouse | Grass | 1,094 | 77 | 10.2 | 35.3 |
| Northern pocket gopher | General | 1,384 | 35 | 7.0 | 62.9 |
| Northern grasshopper mouse | General | 1,372 | 22 | 2.4 | 40.3 |
| Richardson's ground squirrel | Grass | 1,384 | 10 | n/a | n/a |
| Long-tailed vole | General | 265 | 1 | n/a | n/a |
| Ord's kangaroo rat | Grass | 263 | -1 | n/a | n/a |
| Prairie vole | Grass | 654 | -1 | n/a | n/a |
| Meadow vole | Grass | 1,384 | -256 | n/a | n/a |

**Table 4.** Summary statistics for generalized additive models of the influence of habitat loss and fragmentation on rodent species abundances across the Canadian Prairies. Predictors include habitat loss (LOSS), grassland edge density (ED), average grassland patch size (AREA), average grassland patch shape complexity (PARA), grassland patch density (PD), and grassland patch cohesion (COH). ED, AREA, PARA, PD, and COH were included in models as interactions with LOSS to account for the dependence of fragmentation effects on the presence of habitat loss. Bolded predictors are significant at p < 0.001 as identified by the F-statistic (F) and estimated degrees of freedom (EDF). These models form the analytical basis for Figures 2–9.

| Species | Predictor | Main Effects |  |  | Effect Mediated by LOSS |  |  |
| --- | --- | --- | --- | --- | --- | --- | --- |
|  |  | edf | F | p value | edf | F | p value |
| Deer mouse | LOSS | **2.4** | **2.5** | **0.0005** | n/a | n/a | n/a |
|  | ED | **3.7** | **6.5** | **$3.4 \times 10^{-7}$** | 5.7 | 0.8 | 0.01 |
| | AREA | 2.5 | 1.4 | 0.03 | $5.5 \times 10^{-7}$ | 0.0 | 0.6 |
|  | PARA | **2.2** | **3.5** | **0.0001** | 5.4 | 1.1 | 0.0002 |
|  | PD | 1.9 | 1.1 | 0.04 | **7.5** | **2.1** | **$4.0 \times 10^{-5}$** |
|  | COH | **1.6** | **3.9** | **$6.8 \times 10^{-6}$** | **3.0** | **1.2** | **$9.3 \times 10^{-5}$** |
| Sagebrush vole | LOSS | $1.2 \times 10^{-6}$ | 0.0 | 0.4 | n/a | n/a | n/a |
| | ED | $8.1 \times 10^{-7}$ | 0.0 | 0.6 | 0.005 | 0.0 | 0.3 |
|  | AREA | 1.8x10<sup>-7</sup> | 0.0 | 0.8 | **2.0** | **2.4** | **6.9x10<sup>-10</sup>** |
|  | PARA | 1.5x10<sup>-7</sup> | 0.0 | 1.0 | 1.0 | 0.2 | 0.06 |
|  | PD | 8.1x10<sup>-7</sup> | 0.0 | 1.0 | 1.6 | 0.4 | 0.02 |
|  | COH | **2.0** | **30.0** | **2.0x10<sup>-16</sup>** | 3.0 | 0.5 | 0.05 |
| Northern grasshopper mouse | LOSS | 0.0003 | 0.0 | 0.6 | n/a | n/a | n/a |
|  | ED | 0.0002 | 0.0 | 0.9 | 0.0002 | 0.0 | 0.6 |
|  | AREA | 0.0001 | 0.0 | 0.5 | **1.0** | **19.2** | **3.3x10<sup>-7</sup>** |
|  | PARA | 0.7 | 1.9 | 0.06 | 0.001 | 0.001 | 0.4 |
|  | PD | **1.5** | **8.4** | **0.0008** | 0.9 | 1.7 | 0.1 |
|  | COH | 0.0001 | 0.0 | 1.0 | 2.1 | 4.5 | 0.07 |
| Olive-backed pocket mouse | LOSS | **2.9** | **43.2** | **1.0x10<sup>12</sup>** | n/a | n/a | n/a |
|  | ED | 0.2 | 0.3 | 0.2 | **1.5** | **19.2** | **1.9x10<sup>-7</sup>** |
|  | AREA | 0.0002 | 0.0 | 0.8 | 0.0004 | 0.0 | 0.6 |
|  | PARA | 0.0002 | 0.0 | 0.8 | 0.6 | 1.3 | 0.09 |
|  | PD | 0.0002 | 0.0 | 0.7 | **4.3** | **18.9** | **0.0001** |
|  | COH | 0.0003 | 0.0 | 0.7 | 2.2 | 10.0 | 0.001 |
| Northern pocket gopher | LOSS | **1.7** | **13.2** | **5.8x10<sup>-5</sup>** | n/a | n/a | n/a |
|  | ED | 6.9x10<sup>-5</sup> | 0.0 | 0.7 | 0.7 | 1.6 | 0.08 |
|  | AREA | 8.0x10<sup>-5</sup> | 0.0 | 0.5 | 0.0001 | 0.0 | 0.5 |
|  | PARA | 7.3x10<sup>-5</sup> | 0.0 | 0.9 | 0.0001 | 0.0 | 0.9 |
|  | PD | 5.4x10<sup>-5</sup> | 0.0 | 0.8 | **4.2** | **14.4** | **0.0008** |
|  | COH | **0.9** | **8.1** | **0.001** | 0.0001 | 0.0 | 0.8 |
| Thirteen-lined ground squirrel | LOSS | 0.09 | 0.1 | 0.1 | n/a | n/a | n/a |
|  | ED | **2.4** | **34.1** | **1.9x10<sup>-12</sup>** | 6.8x10<sup>-5</sup> | 0.0 | 0.8 |
|  | AREA | 5.4x10<sup>-5</sup> | 0.0 | 1.0 | **2.8** | **12.3** | **1.4x10<sup>-5</sup>** |
|  | PARA | **2.9** | **16.5** | **2.5x10<sup>-5</sup>** | 1.8 | 5.4 | 0.01 |
|  | PD | 1.6x10<sup>-5</sup> | 0.0 | 0.6 | **9.6** | **55.5** | **1.2x10<sup>-13</sup>** |
|  | COH | **0.9** | **9.8** | **3.0x10<sup>-5</sup>** | 2.5 | 4.7 | 0.02 |

All three species that responded to habitat loss (LOSS), estimated here as the summed proportions of annual cropland, perennial livestock forage, and urban development, did so up to a threshold of ~40% LOSS, above which the trajectory of their response changed. Olive-backed pocket mice, considered an arid-adapted grassland obligate, responded negatively to increasing habitat loss to up to 70% LOSS, above which their response reversed to positive (Fig. 2a). The other two grassland obligates (i.e., sagebrush voles and thirteen-lined ground squirrels) showed no response to habitat loss. Predictably for a habitat generalist, deer mice responded positively to <50% LOSS and >90% LOSS but reversed its response to negative where 50-90% LOSS occurred (Fig. 2b). Northern pocket gophers, another habitat generalist, responded positively to <10% LOSS, negatively between 10-80% LOSS, and positively to >90% LOSS (Fig. 2c).

**Fig. 2.**
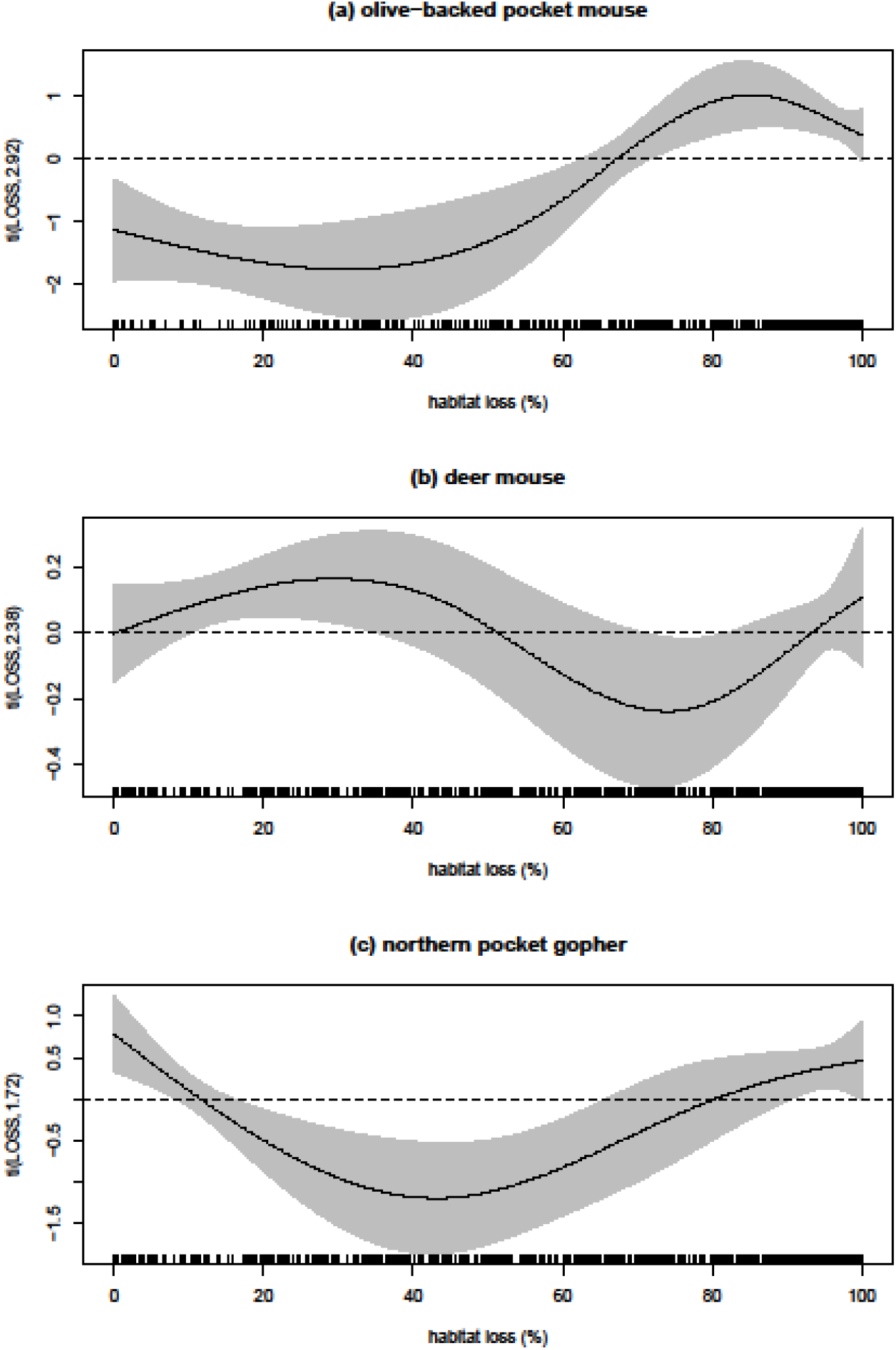
Estimated effect of habitat loss (%; including cropland, perennial livestock forage, and urban and industrial land use types) on **(a)** olive-backed pocket mouse, **(b)** deer mouse, and **(c)** northern pocket gopher abundances across the Canadian Prairies. Trends were modeled on the scale of the linear predictor (solid black line; dotted black line indicates no effect) with 95% confidence intervals (grey shade).

Half of the rodent species considered here responded consistently with predictions based on their specificity for grassland habitat. Thirteen-lined ground squirrels were the only obligate species to respond predictably to grassland fragmentation, increasing linearly with increasing cohesion, or connectivity among patches (Fig. 3a). Thirteen-lined ground squirrels also responded predictably to grassland edge density, declining in abundance to up to 50 m/ha (Fig. 3b). In contrast, two habitat generalists responded as predicted to at least one metric of grassland fragmentation. Northern grasshopper mice responded positively to decreasing average grassland patch size, showing elevated abundances where patches were <1 ha in landscapes with <40% or >90% LOSS (Fig. 4a). And northern pocket gophers responded positively to landscapes with 2-4 grassland patches/100 ha in landscapes with 20-60% LOSS (Fig. 4b).

**Fig. 3.**
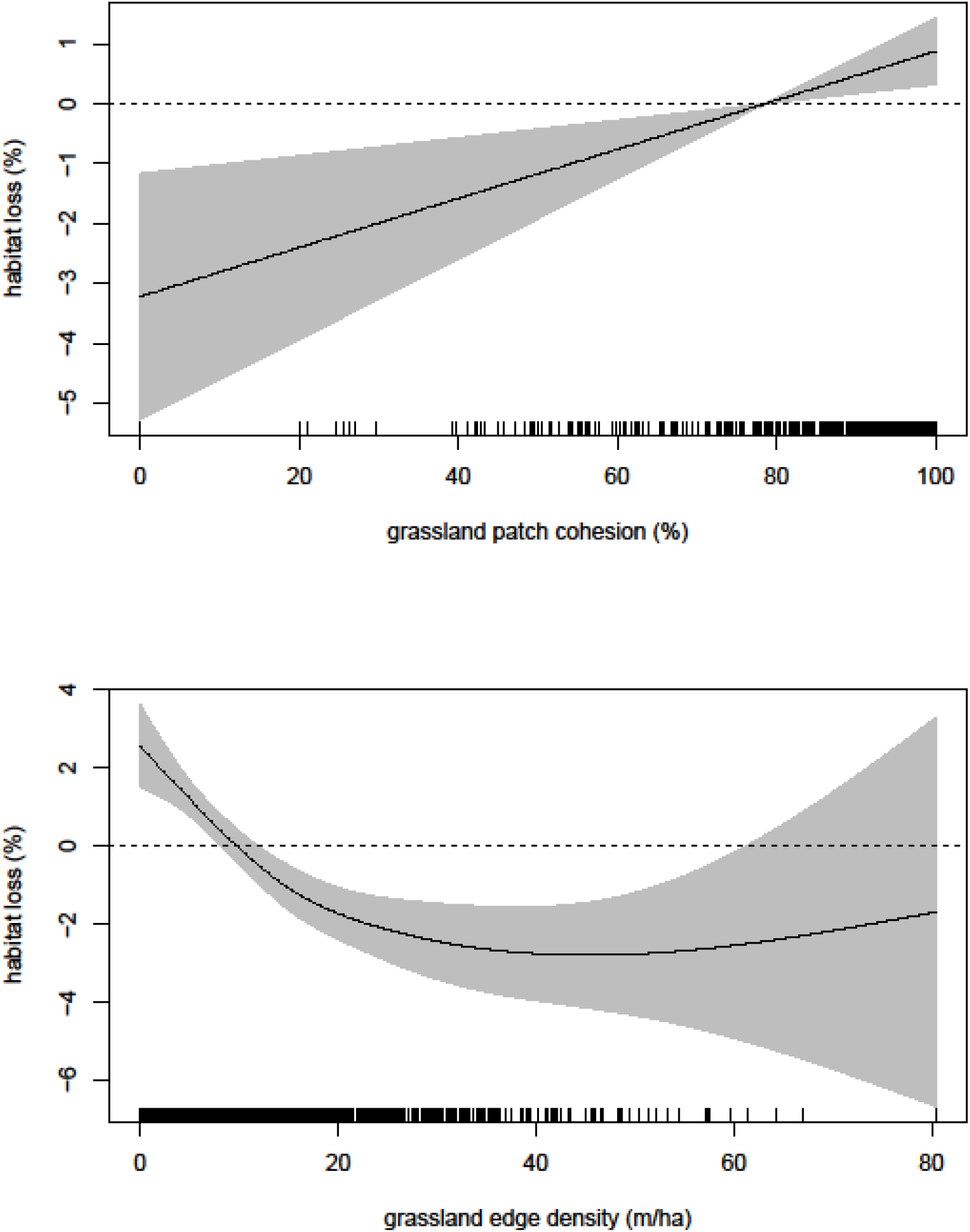
Responses to fragmentation effects that were consistent with predictions based on habitat specificity, including the estimated effect of **(a)** grassland patch cohesion (%) and **(b)** grassland edge density (m/ha) on thirteen-lined ground squirrel abundances across the Canadian Prairies. Trends were modeled on the scale of the linear predictor (solid black line; dotted black line indicates no effect) with 95% confidence intervals (grey shade).

**Fig. 4.**
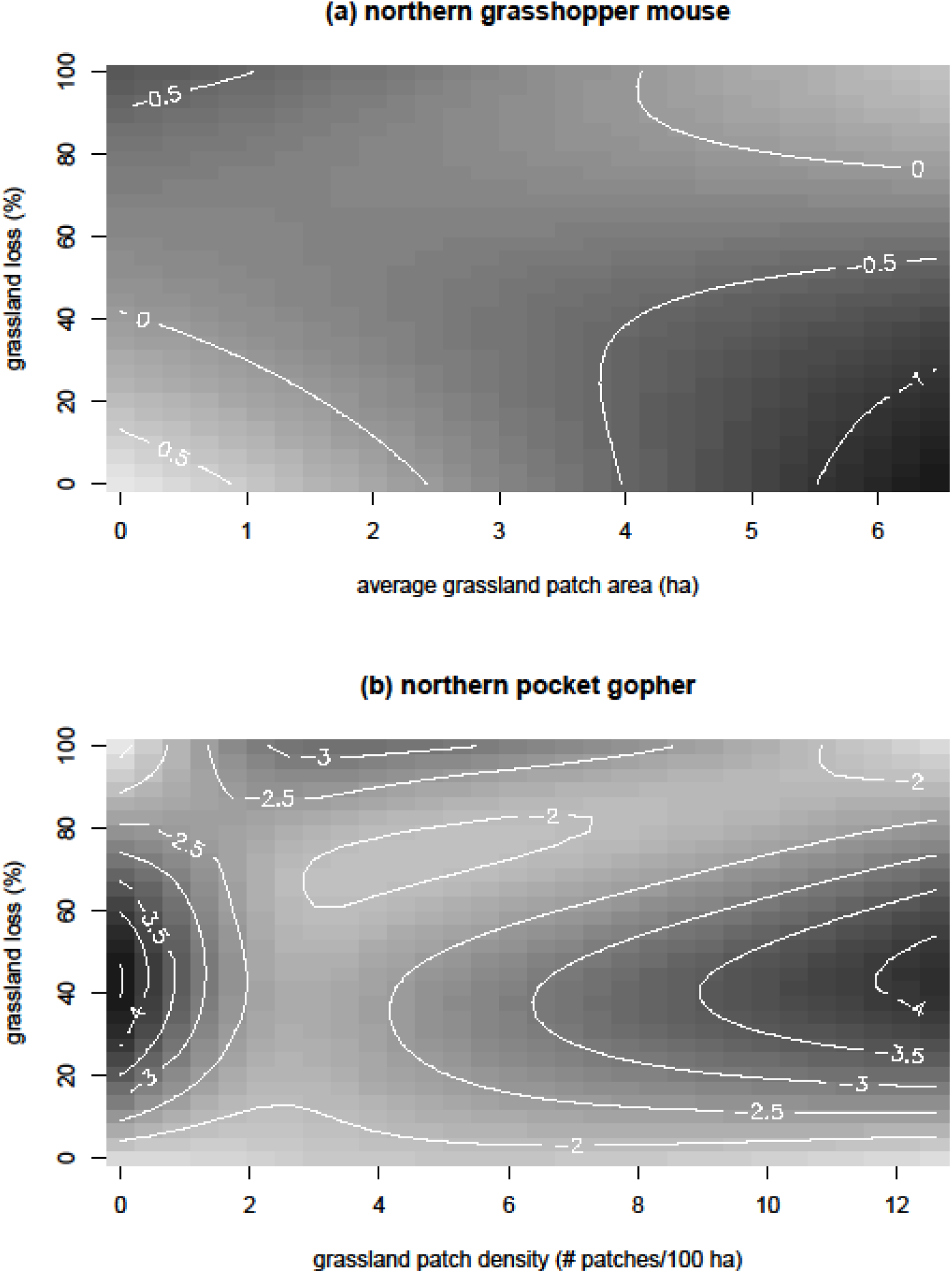
Responses to fragmentation effects independent of habitat loss that were consistent with predictions based on habitat specificity, including the estimated effect of **(a)** average grassland patch area (ha) on northern grasshopper mouse abundance and **(b)** grassland patch density (# patches/100 ha) on northern pocket gopher abundance across the Canadian Prairies. Trends were modeled on the scale of the linear predictor (white contour lines; positive effects shaded light, negative effects shaded dark).

Responses to fragmentation that were inconsistent with our predictions were also observed among grassland obligates. Sagebrush voles declined in abundance with increasing grassland patch cohesion (Fig. 5a), and in 30-80% LOSS shifted from a positive response where average grassland patch size was <2 ha to a negative response where patches were >5 ha in size (Fig. 6a). Thirteen-lined ground squirrels responded positively to landscapes containing >5 patches/100 ha and 20-80% LOSS (Fig. 6b). Olive-backed pocket mice responded positively to landscapes with grassland edge densities >50 m/ha and 30-80% LOSS (Fig. 6c). Responses from habitat generalists were also inconsistent with our predictions. Deer mice responded positively to grassland patch cohesion (Fig. 5b), negatively to >50 m/ha of grassland edge (Fig. 5c), and negatively to simple grassland patch shapes (i.e., <0.06 PARA; Fig. 5d). Northern grasshopper mice responded negatively to increasing grassland patch density (Fig. 5e). Northern pocket gophers responded negatively to landscapes containing >4 patches/100 ha and 20-80% LOSS (Fig. 6d).

**Fig. 5.**
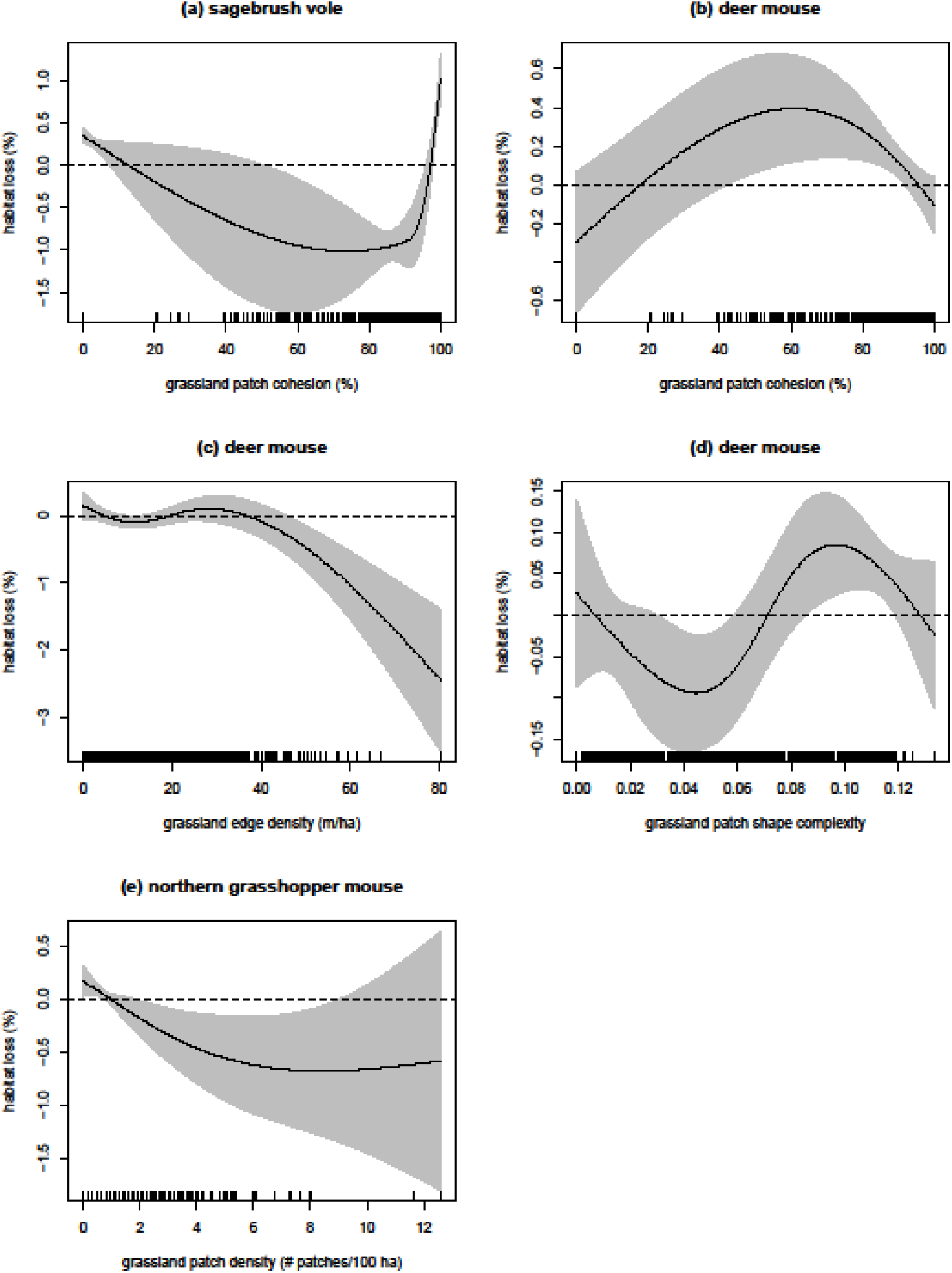
Responses to fragmentation effects that were inconsistent with predictions based on habitat specificity, including the estimated effect of **(a)** grassland patch cohesion (%) on sagebrush vole abundance, the effects of **(b)** grassland patch cohesion, **(c)** grassland edge density (m/ha), and **(d)** grassland patch shape complexity on deer mouse abundance, and the effects of **(e)** grassland patch density (# patches/100 ha) on northern grasshopper mouse abundance. Trends were modeled on the scale of the linear predictor (white contour lines; positive effects shaded light, negative effects shaded dark).

**Fig. 6.**
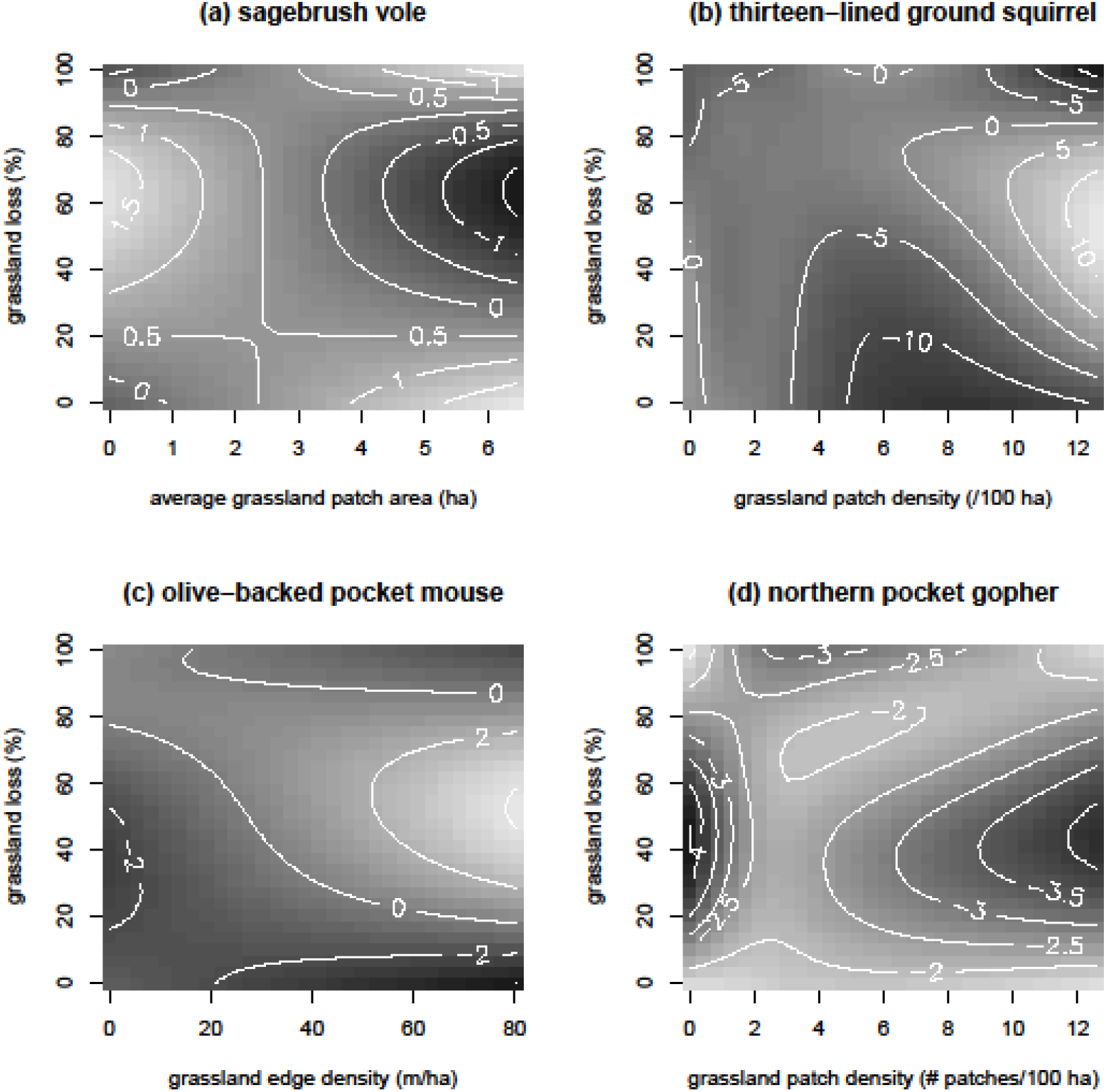
Responses to fragmentation effects independent of habitat loss that were inconsistent with predictions based on habitat specificity, including the estimated effect of **(a)** average grassland patch area (ha) on sagebrush vole abundance, **(b)** grassland patch density (# patches/100 ha) on thirteen-lined ground squirrel abundance, **(c)** grassland edge density (m/ha) on olive-backed pocket mouse abundance, and **(d)** grassland patch density (# patches/100 ha) on northern pocket gopher abundance. Trends were modeled on the scale of the linear predictor (white contour lines; positive effects shaded light, negative effects shaded dark).

Responses were also observed where fragmentation metrics characterized habitat heterogeneity instead of fragmentation per se. These responses were observed among species that responded to fragmentation effects independent of habitat loss where little to no habitat loss had occurred (i.e., <20% LOSS). In these landscapes, sagebrush voles responded positively to grassland patches that were >4 ha in size (Fig. 7a). Olive-backed pocket mice responded negatively to increasing grassland patch density (Fig. 7b). Thirteen-lined ground squirrels showed decreased abundance with increasing grassland patch density (Fig. 7c). Deer mice responded positively to landscapes containing 4-12 grassland patches/100 ha (Fig. 7d).

**Fig. 7.**
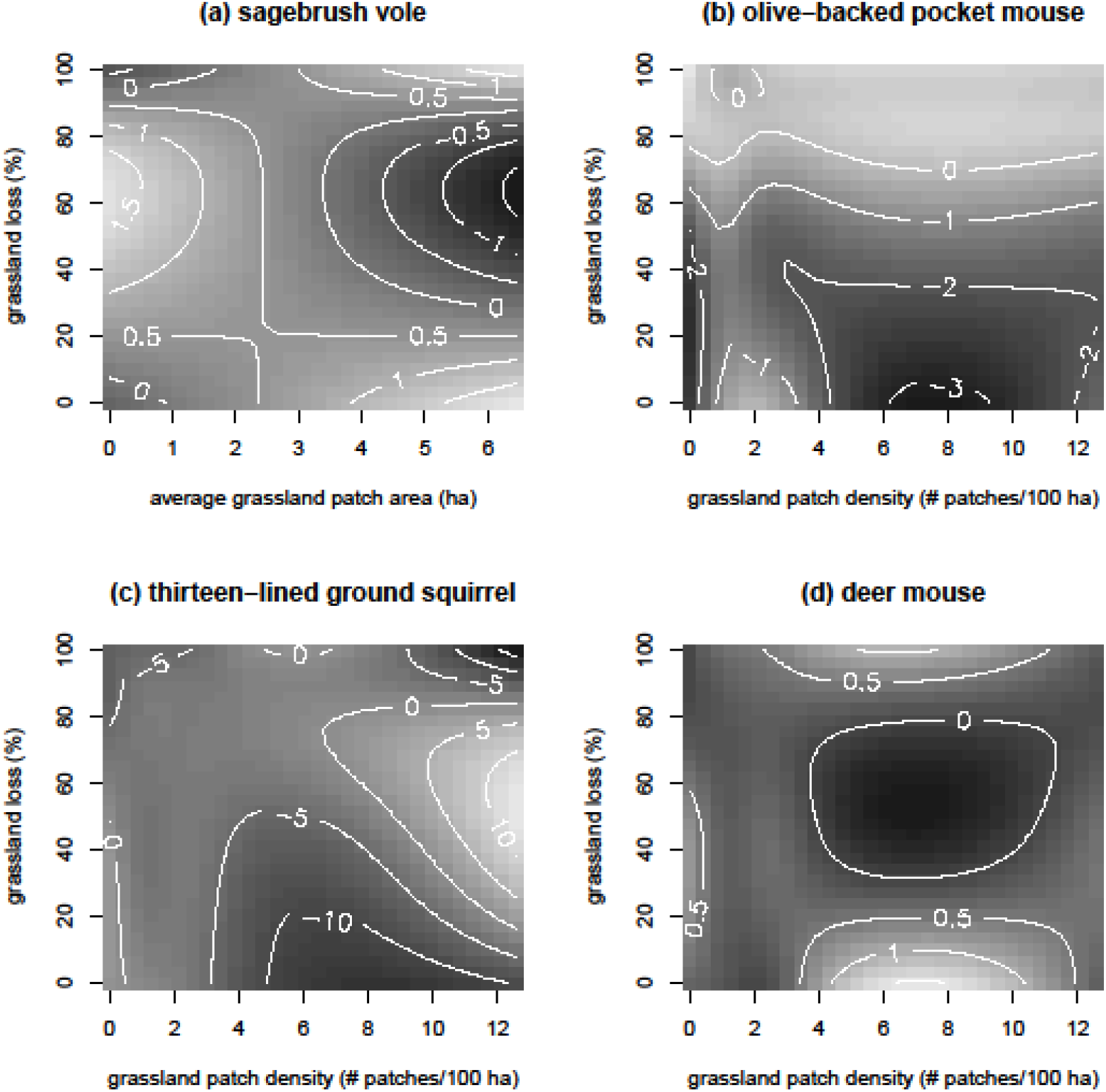
Responses to habitat heterogeneity in landscapes where <20% habitat loss has occurred, including the estimated effect of **(a)** average grassland patch area (ha) on sagebrush vole abundance and grassland patch density (# patches/100 ha) on **(b)** olive-backed pocket mouse abundance, **(c)** thirteen-lined ground squirrel abundance, and **(d)** deer mouse abundance. Trends were modeled on the scale of the linear predictor (white contour lines; positive effects shaded light, negative effects shaded dark).

## Discussion

We found rodent populations showed stronger responses to changes in habitat composition than habitat configuration on the Canadian Prairies. Our results found the combined effects of habitat loss and fragmentation explained an average 9x less variation in rodent abundances relative to habitat composition, soil texture, and climate variation within the landscape. Additionally, only three species responded to habitat loss (i.e., olive-backed pocket mice, deer mice, and northern pocket gophers) while all species responded to at least one metric of habitat fragmentation. The only two studies comparable to this one found lower rodent diversity in fragmented landscapes (Massa et al. 2013) and shifts in assemblage composition with increasing habitat fragmentation (Pena et al. 2003). Studies not including fragmentation effects per se also found strong associations between rodent dynamics and habitat composition at the landscape-level (e.g., Heisler et al. 2013; Rodriguez and Peris 2007; Torre et al. 2015). However, too few fragmentation studies on rodent assemblages have been conducted at the landscape level to draw definitive conclusions. But on the Canadian Prairies, habitat loss and the corresponding change in habitat configuration appear to play minor roles in delineating rodent population responses to landscape-level habitat modification for agriculture.

This is the first study to distinguish fragmentation effects from those of habitat loss on rodent assemblages at a landscape level. We found habitat specificity does predict population responses to habitat loss and fragmentation among grassland obligates and generalists alike, providing further evidence that modifying natural habitats for agricultural production has potential to threaten these obligate species with regional extirpation as landscapes are simplified in habitat composition, shifting rodent assemblages towards generalist-dominated communities (Heisler et al. 2013). In fact, we also found half of the species considered here (i.e., olive-backed pocket mice, deer mice, and northern pocket gophers) reversed the trajectory of their responses to land modification in landscapes where ~50% of the natural habitat was converted, suggesting a threshold above which land modification begins homogenizing habitat heterogeneity with similar consequences for biodiversity. This threat persists on the Canadian Prairies as remaining habitat is converted to up to an average 3.5% per year (Gage et al. 2016). Reduced rodent diversity in heavily modified landscapes has implications for the ecosystem functions they facilitate among higher trophic levels. For example, meadow vole irruptions in this study area are now dampened and less consistent (Poulin et al. 2001; Heisler et al. 2014), with potential to influence the persistence of some prairie predators, including the endangered burrowing owl (Todd et al. 2001) and the short-eared owl, a species of special concern (Colvin and Spaulding 1983). Habitat generalists such as deer mice and northern grasshopper mice appear to benefit from land modification, potentially increasing the prevalence of zoonotic diseases in populated rural areas (Mills and Childs 1998; Ostfeld and Holt 2004; Mills 2005; Hjelle and Torres 2010; Jonsson et al. 2010). Further research on how landscape-level population responses inform the spatial distributions of these rodent species and affect their functional roles in highly modified landscapes is needed in this study area and elsewhere.

Several species responded to fragmentation metrics characterizing habitat heterogeneity in landscapes where grasslands remain intact (i.e., 0-20% LOSS). In these landscapes, fragmentation metrics estimate heterogeneity in grassland patch configuration amongst a mosaic of other natural habitats (i.e., wetlands, shrubland, aspen bluffs, forest patches; Li and Reynolds 1995) instead of fragmentation per se (Andren et al. 1994; Hanski and Ovaskeinan 2002; Fahrig 2003; Betts et al. 2007). For example, both deer mice and thirteen-lined ground squirrels responded positively to increasing heterogeneity in intact landscapes (deer mouse: >4 patches/100 ha; thirteen-lined ground squirrels: patches 2-4 ha in size in low densities). These and other species also responded positively to increasing grassland fragmentation where considerable habitat loss occurred (i.e., >80% LOSS), identifying similar responses among rodent species between heterogeneous and fragmented landscapes, which has been observed in other studies (e.g., Jonsen and Fahrig 1997; Holland et al. 2004; Thies et al. 2003; Holzschuh et al. 2010). This may be because the structural complexity of habitat composition in heterogeneous landscapes provides a greater diversity of habitat niches through which a higher diversity of rodent species can exploit environmental resources (Bazzaz 1975; Tews et al. 2004; Stein et al. 2014). Several studies of local-level fragmentation effects on biodiversity also observed higher biodiversity in agroecosystems (e.g., Lack 1969; Bazzaz 1975; Allouche et al. 2011). Additionally, Fahrig (2017) found over half of all significant fragmentation effects were positive, suggesting these effects may not be uniformly negative on biodiversity.

Habitat specificity may not be a definitive indicator of sensitivity to habitat loss and fragmentation among rodent species in this study area. None of the rodent species considered here responded consistently to all fragmentation metrics as predicted by their respective specificity towards grassland habitat. All grassland obligates except olive-backed pocket mice failed to decline in abundance with increasing percent habitat loss, and all three grassland obligates responded positively to at least one metric of increasing grassland fragmentation. These anomalies may be due to assuming all habitat obligates are dependent on the availability of a single habitat type. Habitat specificity is a proxy for the trade-off between a species’ ability to exploit a range of resources relative to its capacity to use each resource (i.e., ecological specialization, or the ‘jack of all trades is a master of none’ hypothesis; MacArthur 1972; Clavel et al. 2010). Use of this proxy requires the assumption that the habitat(s) occupied by obligates encompass all their required biotic and abiotic conditions, while unoccupied habitats are missing key conditions for population persistence (DeVictor et al. 2010). Some taxonomic groups exhibit highly ordered responses to habitat loss according to each species’ position along habitat specificity gradients (Patterson and Atmar 1986; Wright et al. 1988; Nupp and Swihart 2000; Presley et al. 2010), identifying habitat specificity as an effective proxy for ecological specialization in some cases. Depending on which aspect of fragmentation is under scrutiny, using habitat specificity as a proxy for susceptibility to deleterious effects of habitat loss and fragmentation may not be informative.

Fragmentation effects inconsistent with predictions of habitat specificity gradients may indicate a mismatch between what constitutes habitat for each species and how habitat is defined in studies. Habitat is one of the most ambiguous terms used in ecology (Hall et al. 1997; Kearney 2006; Bamford and Calver 2014), referring to all biotic and abiotic conditions influencing population persistence of a species (Morris 2003). Comparisons of habitat use among species is facilitated by Hutchinson’s concept of the ecological niche, defined as the n-dimensional hypervolume of resources used, each of which is represented by an axis (i.e., food, water sources, shelter, parturition sites, etc.; Hutchinson 1957). Along each axis, each species displays a wide or narrow tolerance or pattern of use relative to other species. A species can therefore be generalist in the use of some resources but specialist in others (Futuyma and Moreno 1988; Clavel et al. 2010). Within this context, the use of habitat specificity as a proxy for ecological specialization is problematic for some species. Habitat is often used synonymously with natural vegetation cover in fragmentation studies (Almeida-Gomez et al. 2015), which may not encompass the gradients of resource uses most limiting to the species of interest in highly fragmented agroecosystems (Fischer and Lindenmeyer 2007; Presley et al. 2010; Betts et al. 2014). For example, olive-backed pocket mice are thought to be restricted to unmodified grassland habitat throughout their geographic range (Hayward and Killpack 1956; Banfield 1974; Lampe et al. 1974; Wilhelm et al. 1981) but elevated abundances were observed in modified landscapes with high grassland edge densities. Large populations of this species may therefore be limited to grassland vegetation in less modified landscapes but released from that limitation in modified landscapes where individuals may take advantage of increased seed production along grassland-cropland edges. Similarly, northern pocket gophers responded negatively to up to 40% habitat loss despite being characterized as a habitat generalist. This fossorial species may not select for vegetative habitat(s) per se but are limited to those that do not experience intra-annual soil disturbances like annual cropland (Salt 2000). This hypothesis does not explain the increase in abundance of this species to up to 100% habitat loss. Future studies should carefully consider which dimension of the ecological niche defines habitat specificity for all species under study prior to assuming the natural vegetation of the study area appropriately characterizes resource limitation among all species (Franklin et al. 2005; Fischer and Lindenmeyer 2007; Flynn et al. 2009; Betts et al. 2014).

The use of habitat specificity as a proxy for ecological specialization may have significant implications for biodiversity conservation when used to inform land modification policies at a national or international scale. The amount of land modified for global agricultural production is expected to increase 10–50% by 2050 (Tilman et al. 2001; Tilman et al. 2011), most likely in developing countries where the world’s biodiversity hotspots remain (Myers et al. 2000; Green et al. 2005). Two strategies, or a combination of both, have been proposed to conserve these remaining hotspots: 1) promoting agricultural intensification in existing agroecosystems considered low priority for biodiversity conservation to save remaining biodiversity hotspots from land modification (i.e., land sparing approach; Green et al. 2005; Phalan et al. 2011; Tilman et al. 2011), or 2) preventing agricultural intensification by improving crop yields instead of continued land modification, maintaining mosaics of natural habitats interspersed among agricultural land-use types where biodiversity conservation is a priority (i.e., land sharing approach; Foley et al. 2011; Wright et al. 2012; Mendenhall et al. 2014). Habitat specificity is now being used to predict biodiversity responses to different conservation strategies, with little to no empirical research to assess its reliability for use to inform global biodiversity conservation (Green et al. 2005; Cardinale et al. 2012; Baudron and Giller 2013). For example, Phalan et al. (2011) found more bird and tree species showed lower densities in 1 km^2^ plots of modified habitat and advocate for land sparing to conserve large expanses of intact natural habitat elsewhere. In contrast, Wright et al. (2012) found not all species maintain high population densities in natural habitats, but some species are now reliant on agricultural land-use types showing high densities in heavily modified agricultural areas. Both studies fail to recognize those species whose population persistence is limited by biotic or abiotic conditions not defined by vegetation, including grassland habitats and agricultural land-use types. Our study finds species exhibit complex, sometimes contradictory responses to habitat fragmentation dependent on the magnitude of habitat loss that has occurred that may not be predicted by habitat specificity. Use of habitat specificity to predict anthropogenic impacts on biodiversity should be done with caution considering the finality of land modification, or at the very least explicitly define the specialization gradient of interest.

## Acknowledgements

We are grateful to all reviewers for their helpful comments on earlier versions of this manuscript. Support was provided by the Institute of Environmental Change & Society (IECS), Natural Sciences and Engineering Research Council of Canada (NSERC; including Discovery Grant 2014-04032 to GLS), Environment & Climate Change Canada, Royal Saskatchewan Museum, University of Regina, Saskatchewan Innovation & Opportunities, and the Saskatchewan Fish & Wildlife Development Fund.

